# Gap perception in bumblebees

**DOI:** 10.1101/315432

**Authors:** Sridhar Ravi, Olivier Bertrand, Tim Siesenop, Lea-Sophie Manz, Charlotte Doussot, Alex Fisher, Martin Egelhaaf

## Abstract

A number of insects fly over long distances below the natural canopy where the physical environment is highly cluttered consisting of obstacles of varying shape, size and texture. While navigating within such environments animals need to perceive and disambiguate environmental features that might obstruct their flight. The most elemental aspect of aerial navigation through such environments is gap identification and passability evaluation. We used bumblebees to seek insights into the mechanisms used for gap identification when confronted with an obstacle in their flight path and behavioral compensations employed to assess gap properties. Initially, bumblebee foragers were trained to fly though an unobstructed flight tunnel that led to a foraging chamber. After the bees were familiar with this situation, we placed a wall containing a gap that unexpectedly obstructed the flight path on a return trip to the hive. The flight trajectories of the bees as they approached the obstacle wall and traversed the gap were analyzed in order to evaluate their behavior as a function of the distance between the gap and a background wall that was placed behind the gap. Bumblebees initially decelerate when confronted with an unexpected obstacle. Deceleration was first noticed when the obstacle subtended around 35° on the retina but also depended on the properties of the gap. Subsequently the bees gradually traded off their longitudinal velocity to lateral velocity and approached the gap increasing lateral displacements and lateral velocity. Bumblebees shaped their flight trajectory depending on the salience of the gap, in our case, indicated by the optic flow contrast between the region within the gap and on the obstacle, which increases with decreasing distance between the gap and the background wall. As the optic flow contrast decreased the bees spent increasing time moving laterally across the obstacles. During these repeated lateral maneuvers the bees are likely assessing gap geometry and passability.

## Introduction

Even with relatively tiny brains insects display a rich repertoire of behaviors, the operation of many of which still remains unclear. Aerial locomotion below the natural canopy is one such behavior that has received increased attention in the last decade from both biologists and engineers alike (Shyy et al., 2016). Natural flight at the small scale of insects, where the sensory and mechanical constraints are particularly challenging, requires the concerted coordination of their computationally parsimonious sensorimotor system (Dudley, 2002). The spatial environment close to the Earth’s surface consists of a myriad of natural and artificial objects that can vary widely in shape, size and texture. This renders the physical environment to be unpredictable and poses challenges to aerial locomotion. Steady level flight for sustained durations is generally unfeasible in this domain with obstacles constantly coming in the way. Thus, in order to achieve safe transit through such environments flying systems need to be adept at perceiving the environment to identify obstacles and devise alternative flight paths. From a biophysical standpoint, apart from performance limitations based on allometric body size scaling other factors, such as collision avoidance and properties of the physical environment, also influence flight trajectories and overall performance of insects (Crall et al., 2015; Dudley, 2002). Terrestrially bound insects utilize a number of strategies at the sensory and motor level in dealing with cluttered and uneven terrain including active and passive body compliance, gait coordination, and preflexion. A commensurate level of understanding is yet to be arrived at for flying animals.

Unlike during legged locomotion where tactile sensory inputs can augment vision in gaining environmental information, flying insects rely only on the latter for safe passage and path planning. For long distance navigation, flying insects might use, apart from vision, other sensory modalities such as odor and geomagnetic fields (Knaden and Graham, 2016) In order for a flying animal to arrive at its intended destination or ensure safe locomotion, at a basic level, the animal needs to process the obstacles that lie in its path and identify gaps. Obstacle and gap detection may thus be considered the most basic element of flight through clutter. A few recent studies have analysed the response of flying insects in minimally cluttered environments and revealed that when confronted with obstacles with varying spacing, insects such as bumblebees and honeybees choose the larger gap (Baird and Dacke, 2016; Ong et al., 2017). This might seem as an obvious response, yet it highlights the active response of insects in avoiding collisions, which otherwise can result in irreparable damage to body and wings. Baird and Dacke (2016) suggested bumblebees may utilize a simple brightness based strategy in making a choice among the different gaps, i.e. bigger gaps are likely to be brighter than smaller gaps. Though a few experiments have observed insects behaving around individual obstacles and minimally cluttered environments, the mechanisms mediating the elemental process of obstacle and gap perception and the factors that influence the assessment of passability are still unclear.

Unlike during legged locomotion where tactile sensory inputs can augment vision in gaining environmental information, flying insects rely only on the latter for safe passage and path planning. For long distance navigation, flying insects might use, apart from vision, other sensory modalities such as odor and geomagnetic fields (Knaden and Graham, 2016) In order for a flying animal to arrive at its intended destination or ensure safe locomotion, at a basic level, the animal needs to process the obstacles that lie in its path and identify gaps. Obstacle and gap detection may thus be considered the most basic element of flight through clutter. A few recent studies have analysed the response of flying insects in minimally cluttered environments and revealed that when confronted with obstacles with varying spacing, insects such as bumblebees and honeybees choose the larger gap (Baird and Dacke, 2016; Ong et al., 2017). This might seem as an obvious response, yet it highlights the active response of insects in avoiding collisions, which otherwise can result in irreparable damage to body and wings. Baird and Dacke (2016) suggested bumblebees may utilize a simple brightness based strategy in making a choice among the different gaps, i.e. bigger gaps are likely to be brighter than smaller gaps. Though a few experiments have observed insects behaving around individual obstacles and minimally cluttered environments, the mechanisms mediating the elemental process of obstacle and gap perception and the factors that influence the assessment of passability are still unclear.

Especially fast flying animals, such as many insect species, rely on optic flow as the main source of spatial information, i.e. on the continuous stream of retinal image changes induced during self-motion and, thus, is particularly relevant for behavior in cluttered environments (Egelhaaf 2006; Egelhaaf et al., 2012). Optic flow has also been shown to aid in estimating flight distance, flight path centering, identifying foraging locations and many other behaviorally relevant tasks, see (Baird et al., 2013; Kern et al., 2012; Serres and Ruffier, 2017; Serres et al., 2008; Srinivasan, 2015; Srinivasan and Zhang, 1997). Observations of the flight trajectory of insects such as flies and bees has shown that insects actively shape the temporal structure of their visual input by employing prototypical flight maneuvers, especially to separate translational from rotational optic flow and, thus, to facilitate discerning spatial information about the surroundings (Braun et al., 2010; Braun et al., 2012; Egelhaaf et al., 2012b). Rotations that are inevitably required to change flight direction are squeezed into brief saccadic turns with rotation velocities often exceeding 4000deg/s (Hateren and Schilstra, 1999). Between saccadic turns the insect keeps its gaze basically straight for more than 80% of overall flight time to induce purely translational optic flow (Boeddeker et al., 2010; Braun et al., 2012; Dickinson, 2005; Hateren and Schilstra, 1999; Mertes et al., 2015; Schilstra and Hateren, 1999). Since only the translational optic flow contains spatial information, this active flight and gaze strategy is believed to facilitate spatial vision and thus may be particularly relevant for navigation in cluttered terrain (Egelhaaf et al., 2014). However this vision-based strategy relying on the closed action perception loop has not been investigated systematically in cluttered environment and, thus, needs further investigation. Specifically, how does an insect react to unexpected obstacles obstructing its flight path? What flight maneuvers does it perform in order to detect gaps and assess passability?

Here we sought to uncover the mechanisms implemented by flying insects in gap identification and perception. Bumblebees are excellent model organisms because much is known about their flight and navigational performance (Baird and Dacke, 2012; Crall et al., 2014; Mirwan and Kevan, 2013; Osborne et al., 2008; Ravi et al., 2013; Riabinina et al., 2014; Lobecke et al. 2018). We presented unsuspecting bumblebees with an altered environment consisting of a wall obstructing their flight path but containing a gap that prevented direct passage to their goal and observed their behavior as they approached and traversed the gap. We analyzed the flight trajectory of the bees in different distances from the gap and computed key visual metrics such as: angle subtended by the obstacle and gap on the retina, mean optic flow and optic flow contrast, in order to identify factors that influence gap identification and assessment of passability. Our data suggest that bumblebees employ an active gazing flight strategy in enabling the identification of gaps and critical environmental parameters that affects safe passage.

## Materials and Methods

### Experiment Setup

Experiments were conducted with individuals from a *Bombus terrestris* colony that was maintained within the lab. A healthy hive sourced from a commercial breeder (Biobest Group NV, Westerlo, Belgium) was placed within a 0.5x0.5.0.3m mesh enclosure that was covered with dark cloth to simulate the natural underground habitat of the bees. The hive enclosure was connected to a flight tunnel 0.25×0.25×1.5m that lead to a 1×1×0.75m foraging chamber where gravity feeders containing 30%/vol. sucrose solution blended with 1% commercial honey were placed. Connections between the hive enclosure, flight tunnel and foraging chamber were made using 30mm ID and 150mm long flexible silicon tubing. Finely ground pollen was placed directly within the hive and bees were permitted to access sucrose in the foraging chamber ad libitum. Consistent foraging flights by numerous (>20) worker bees were observed within one day of immigration to the enclosure. The bees and hive were given one week for habituation to the environment before experiments.

During experiments gates on either sides of the flight tunnel were used to regulate traffic, and only one bee at a time was permitted to enter the flight tunnel. Only bees returning to the hive were considered for analysis. The experiment procedure will be described from the perspective of the bee returning to the hive as per Fig. 1a. An obstacle was created within the flight tunnel by placing an artificial vertical wall that contained a rectangular hole that was 50mm wide and started from the middle extending to the top, see Fig. 1a. The sidewalls of the tunnel were lined with an achromatic random checkerboard pattern while the floor was lined with a random cloud with spatial frequencies varying by 1/f, similar to the one used by (Monteagudo et al., 2017). A second vertical wall was placed behind the wall containing the gap. The same checkerboard pattern was also placed on both obstructing verticals, Fig. 1. Five different experiment conditions were tested where the distance between the gap and rear wall was varied by 550, 300, 150, 50 or 0 mm. During the different scenarios the wall containing the gap was always placed at 0.9m from the entrance of the tunnel, see Fig. 1a. Twenty flights were recorded for each condition while the conditions were varied pseudo-randomly between each recording. Once the bees approached and passed the gap the rear wall was removed to permit their onward flight back to the hive. For the condition when the rear wall was adjacent to the gap (d = 0mm) passage was obviously impossible, once 20 sec of recording was completed, the wall with the gap and rear wall were both removed by opening the roof of the flight tunnel. We also observed the flight of the bees when the wall behind the gap was lined with non-textured white paper and placed immediately adjacent to the gap, i.e. similar to the d = 0mm, Fig. 1.

**Fig.1.**
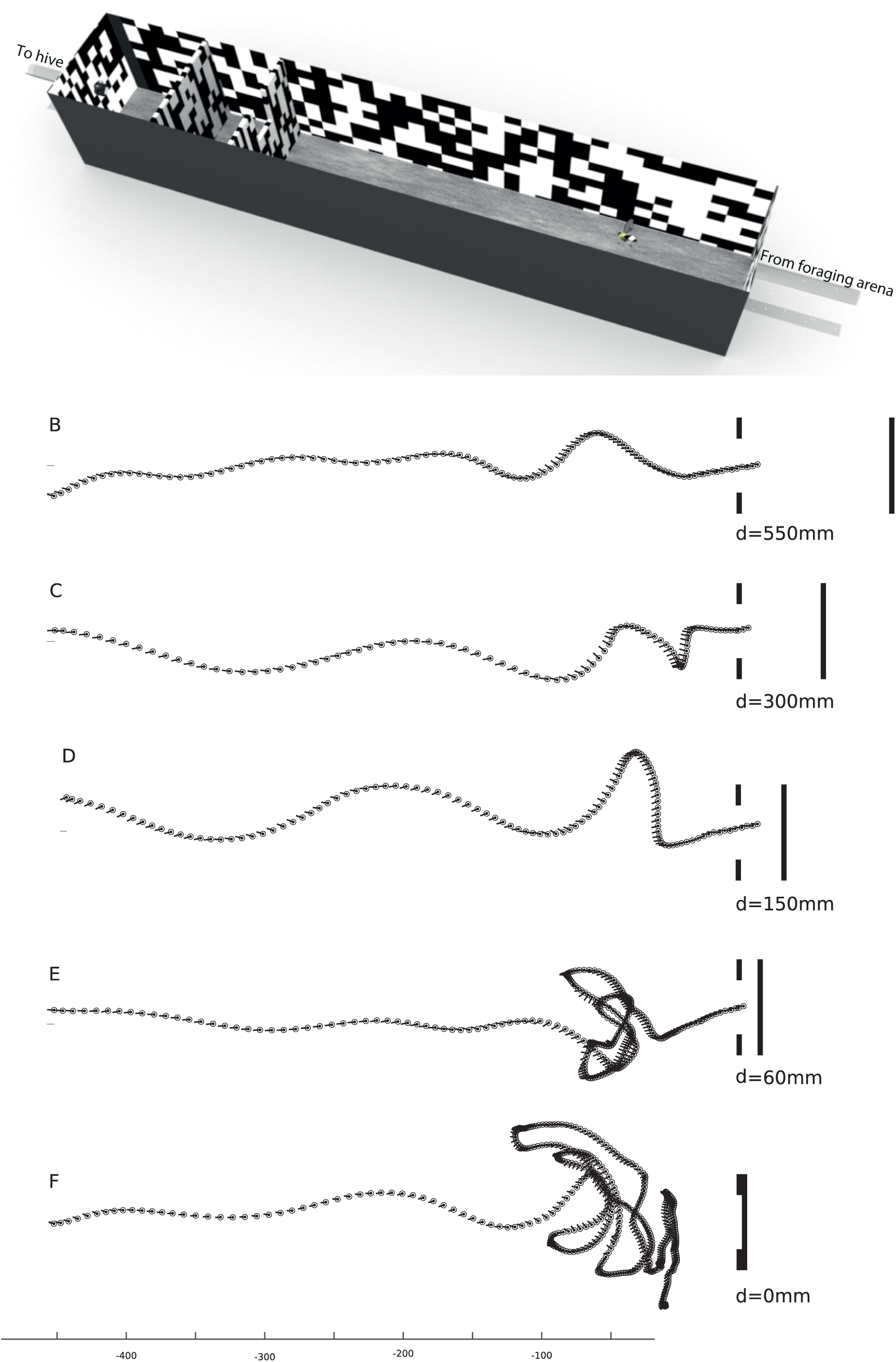
a) Schematic of the experiment setup, d represents the distance between the wall containing the gap to the rear wall. Only the flight of bees returning to the hive from the foraging arena was considered for analysis. Sample flight trajectory of a bee when the distance between the gap and rear wall was 550mm (b), 300mm (c), 150mm (d), 60mm (e) and 0mm (f). The gap and rear wall on the right are for illustrative purposes only and not to scale.

To ensure we captured the response of naïve bees dealing with a complex environment and negotiating a gap, experiment bouts lasted no longer than 1 hour, and the gap and rear wall were removed after each flight recording to inhibit the bees from becoming familiar with the experiment paradigm. Bees were not individually marked in this study, though this increased the possibility of taking unequal numbers of measurements amongst the different individuals for each condition, the likelihood was greatly reduced since consecutive flights were taken from different bees returning to the hive from the foraging arena. Additionally, due to the large number of foragers and flight trajectories recorded it is likely to be representative of the population. All experiments were completed within five consecutive days.

### Fight Trajectory Analysis and Optic Flow Estimation

An Optronis CR6 high-speed camera was placed 1.7m above the midline of the flight tunnel looking directly downward. The flights of the bees were recorded at 200Hz, and a region covering 950mm leading to the gap was kept in the field of view. During post processing lens distortion was corrected by using standard MATLAB Image Processing Toolbox routines. An object of known dimension was placed within the field of view at mid-height of the tunnel and related to the pixels in the rectified image for 2D spatial calibration. Custom MATLAB code was written to process each frame and fit an ellipse to the body of the bees; subsequently the centroid location, body length and heading were all measured over the entire flight. The bees displayed a wide diversity in flight behaviors. In the flight tunnel flights ranged from appearing to explore the space to making directed flights along the tunnel. Only flights of individuals that appeared to be returning from foraging trips, considered as those bees that made a steady and direct flight towards the gap, were used for analysis. At least one such flight was observed every minute. Among all the flights recorded the body length of the individual bee varied by less than 5% indicative of the nominally constant altitude maintained during the entire flight. In order to attenuate digitizing error the flight trajectories were passed through a 30Hz 2^nd^ order Butterworth filter. Flight speed along the longitudinal and lateral directions was estimated by differentiating the flight trajectory along the respective axis and applying a coordinate transformation matrix to obtain body centered values. Heading orientation was calculated with respect to the flight tunnel using the right-hand-rule. Since the flights were recorded only through a single perspective, pitch and roll could not be measured.

Geometric optic flow measured as the angular displacement of the vector between an arbitrary point in space and the retina due to relative motion, see EQU1 was calculated in MATLAB using the flight trajectory and flight tunnel geometry. Here, for each flight in all conditions the true optic flow of only the wall containing the gap and the rear wall was calculated using the respective flight trajectory, assuming constant head-body alignment, a spherical eye and the retina approximated as a point. A similar approach has been implemented in numerous previous studies (Bertrand et al., 2015; Floreano et al., 2010; Serres and Ruffier, 2017; van Breugel et al., 2014).

A total of 100 flights (20/condition x 5 conditions) were recorded, and statistical significance of the variation in quantities between experiment conditions was tested using a on way ANOVA and a Tukey Post-Hoc test confirmed significant conditions within the group. For comparison of quantities within each experiment condition a paired t-test was used to assess statistical significance. No statistical analysis was conducted for the condition where a glossy white background was placed immediately in the rear of the gap, only five flights was permitted and the analysis was qualitative.

## Results

Upon entering the flight tunnel all bees took off and flew smoothly as they approached the unfamiliar wall blocking their flight path. For all experiment conditions enroute to the gap the flight trajectories of the bees were not straight, but contained some smooth lateral movements, see Fig. 1b-f. The bees performed increased lateral maneuvers closer to the gap as the distance between the gap and rear wall was reduced, see Fig. 1d-f. For conditions when the distance between the gap and rear was <60mm the bees engaged in forward facing crescent-shaped maneuvers close to the gap (<100mm), see Fig. 1e, prior to passing through. When the gap was not present i.e. the rear wall was directly adjacent to the gap, the bees continued to preform crescent flight while facing forwards close to the center of the tunnel with increasing arc size. None of the bees attempted to pass through in this condition, see Fig. 1f. This also included the condition when a non-textured white wall was placed immediately adjacent to the gap, similar to the d = 0mm condition (data not shown). An apparent increase in the sideward component in the flight path is evident with decreasing distance between the gap and rear wall. These crescent shaped flight paths bear nominal similarity to learning flights of bumblebees after they leave their nest hole and are assumed to gather information about its surroundings (e.g. Lobecke et al., 2018; Philippides et al., 2013).

In order to quantify the flight trajectories, the tunnel was binned into six segments and the longitudinal flight speed of the bees within each bin was calculated, see Fig. 2a. Irrespective of the distance between the gap and rear wall, the flight speed among the different individuals remained statistically similar when they were >375mm to the gap, Fig. 2a (F(4,95) = 0.79, p = 0.55). The flight speed of the bees within this region was nominally similar to those reported by (Baird et al., 2010) where a similar sized tunnel was used. However there exists considerable variation in flight speed (up to 1.5x variation in magnitude) among the different flights across all conditions. At distances <375mm the bees approached the gap while steadily decelerating wherein the rate of deceleration was dependent on the distance between the gap and background, Fig. 2a.

**Figure 2.**
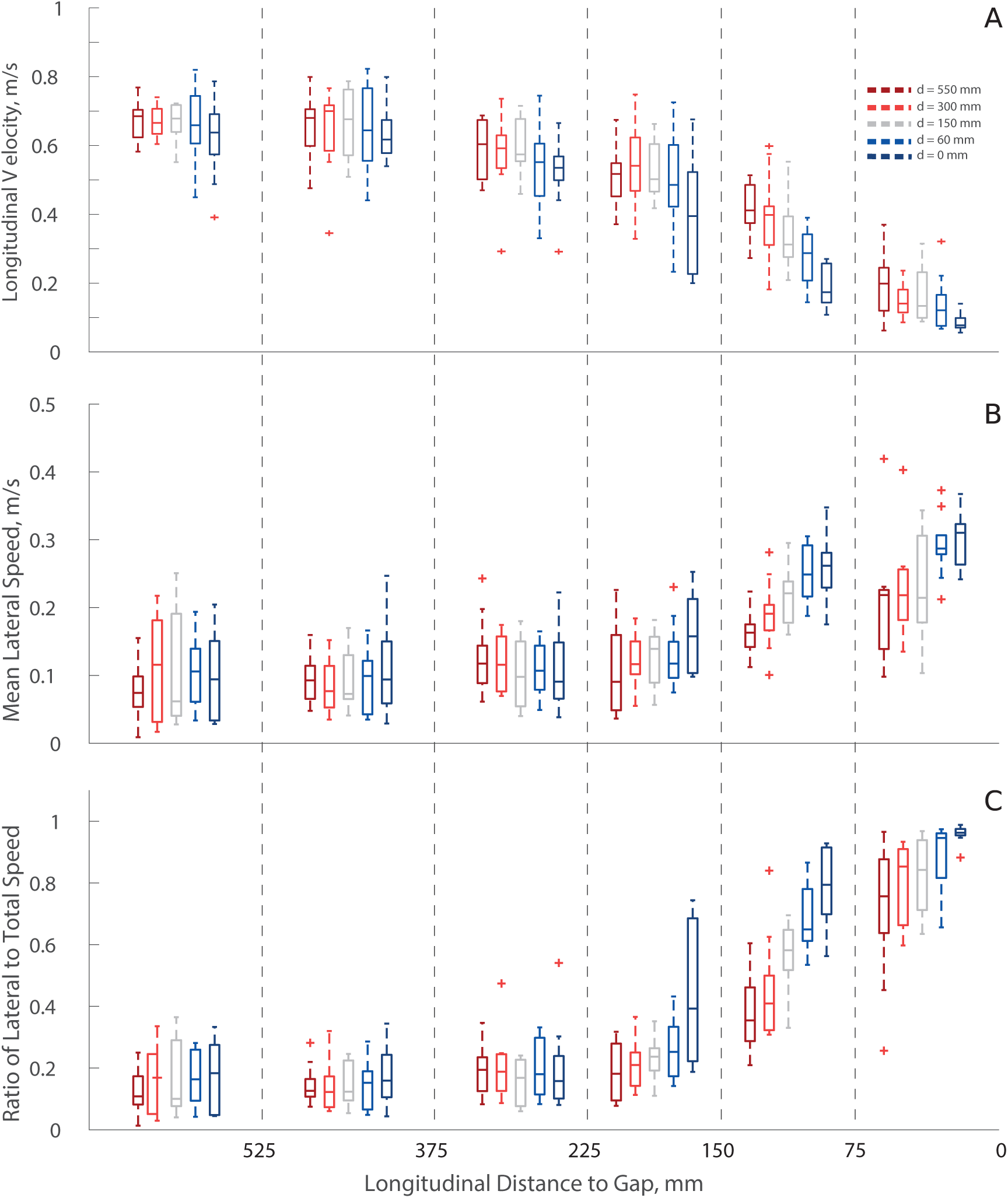
The flight tunnel was binned into six sections leading to the gap. a) The absolute mean longitudinal velocity of the bees at different longitudinal distances from the gap. b) The absolute mean lateral velocity of the bees at different sections of the flight tunnel. c) Ratio of the mean absolute lateral velocity and the total velocity.

The mean longitudinal velocity of the bees was significantly lower when the distance between the gap and rear wall was <150mm (d = 150mm p = 0.043, d = 60mm p = 0.029 & d = 0mm p = 0.0068). When d > 150mm the longitudinal velocity was lower but not statistically significant, d = 300mm p =0.068 & d = 550mm p = 0.075. For the d = 550, 300 and 150mm the flight speed of the bees was significant reduced when they were <225mm to the gap compared to when they were <375mm (d = 550mm p = 0.022, d = 300mm p = 0.046 and d = 150mm p = 0.013), Fig. 2a. When the bees where <150mm to the gap there was a monotonic reduction in their mean speed with decreasing distances between the gap and rear wall (F(4,95) = 3.18, p = 0.02).

For all experiment conditions, the mean absolute lateral speed of the bees was small but existent at large distances to the gap (>225mm) and monotonically increased as they approached the gap, see Fig. 2b. Unlike the longitudinal speed, the mean absolute lateral flight speed did not become significantly different across the different conditions until the bees were <225mm; in this condition the bees’ lateral velocity was maximum when the rear wall was adjacent to the gap (d = 0 p = 0.00935), Fig. 2b. Similar to the forward speed, the rate of increases in the mean lateral speed of the bees was also dependent on the distance between the gap and rear wall. By comparing the ratio of mean lateral speed to the total speed of the bees it is evident that the bees summarily increase their mean lateral movement over longitudinal movement, Fig. 2c. For the non-passable condition where d = 0 over the duration of the recording the bees mostly moved laterally at distances <75mm to the gap (Fig. 2b&c). The lateral distance traveled by the bees within each segment also increased, as they got closer to the gap. This can also be qualitatively observed in the representative flight trajectories, Fig. 1b-f.

To further understand the mechanics of the lateral movements and the flight maneuvers performed close to the gap for conditions where the distance between gap and rear wall was small, the total acceleration of the bees in the body coordinate system at three different segments along the tunnel was represented as a rose histogram, Fig. 3. The length of each angular column in the rose histogram indicates probability of the total acceleration of the bee to be within the range of the respective bin. This was done for the d = 0mm condition where lateral movements were most significant. The angle between the total acceleration and the long axis of the body was binned into twenty segments of 18° width for all recorded flights. When the bees were >375mm from the gap, the total acceleration was oriented laterally with respect to their body and minimally in longitudinal direction. Between 375 – 150mm to the gap the bees began decelerating (Fig. 3b), and this is evident in the acceleration histogram where the total acceleration was distributed mostly laterally with a rearward skew (2^nd^ & 3^rd^ quadrant). In the proximal regions of the gap (<150mm) the total acceleration was predominantly orientated orthogonal to the long body axis, Fig. 3c. The magnitude of acceleration, as indicated by the hue in Fig. 3, was also lowest when the bees were far from the gap. It progressively increased near the gap where the flight was characterized by the largest accelerations oriented nominally orthogonal to the longitudinal axis, Fig. 3a-c.

**Figure 3.**
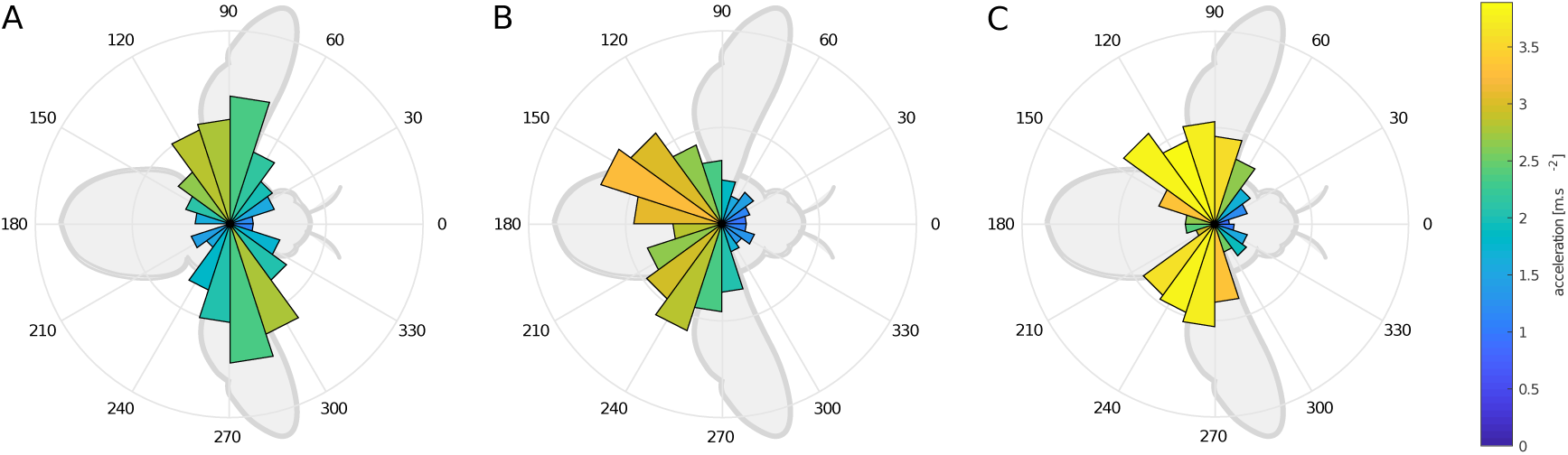
Rose histogram of the total acceleration with respect to long body axis among all flights at different section of the flight tunnel a) 300mm -200mm, b) 200mm – 100mm & c) 100mm – 0mm. The length of each angular column in the rose histogram indicates the probability of the total acceleration of the bee to be within the range of the respective bin. The hue represents the magnitude of mean acceleration within the respective angular bins.

### Optic Flow Analysis

The above results clearly revealed that flight behavior of bumblebees is strongly affected by the distance between the gap and the rear wall. The most likely cue providing information about the spatial layout under the different conditions is the optic flow within the gap and in the adjacent parts of the visual field. Therefore, we determined the optic flow difference between inside and outside the gap. The mean of the absolute difference in total geometric optic flow across the inside and outside edge of the gap (±12 mm along gap edge) is presented Fig. 4a. As expected, the difference in optic flow across the edge of the gap was low when the bees were far from the gap and it progressively increased as the bees neared the gap. Until the bees were <225mm from the gap the difference in mean optic flow across the edge decreased significantly with decreasing distance between the gap and the rear wall. However, in the near vicinity of the gap (<150mm) the mean optic flow difference across the edge of the gap was relatively lower and not statistical significant across the d = 550 – 60mm conditions (F(4,95) = 0.28, p = 0.74), Fig. 4a. The optic flow difference for the d = 0 is nonzero (Fig. 4a) because of the offset in the wall position due to their thickness (1.5mm). The mean optic flow on the wall containing the gap and the rear wall when the bees were <150mm to the gap for all conditions is presented in Fig. 4b-f. A sharp discontinuity in the mean optic flow at the edge of the gap is present when the rear wall is further away from the gap, consequently, the distinctness of the gap clearly decreases with decreasing distance between the gap and the rear wall in the different experiment conditions.

**Figure 4.**
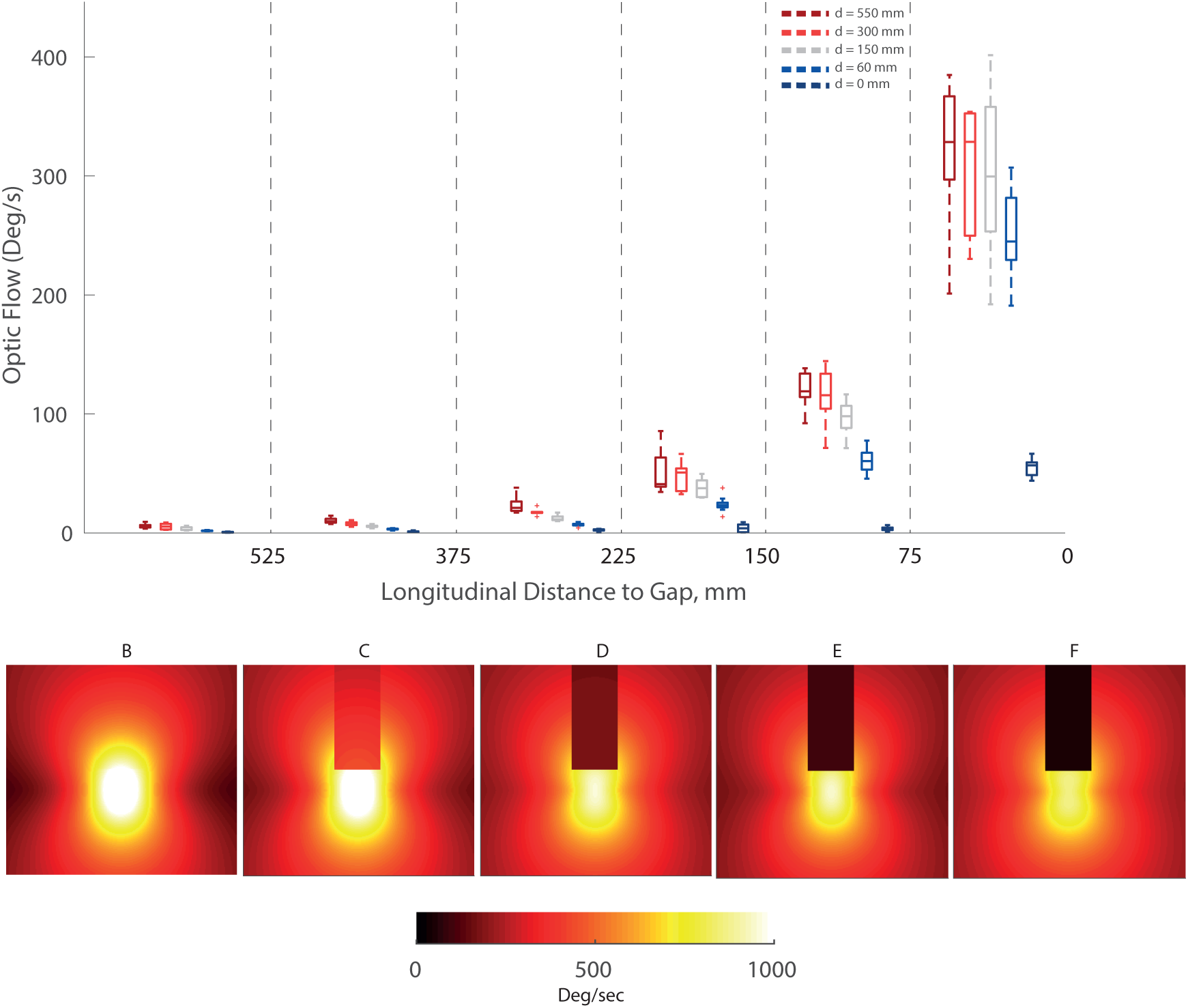
a) The mean of the absolute difference in total optic flow across the inside and outside edge of the gap (±12 mm along gap edge) at different sections of the flight tunnel. b-f) Heat map showing the mean geometric optic flow over the wall containing the gap across all flight trajectories when the bees were <75mm to the gap. The geometric optic flow was calculated taking into account both the longitudinal and lateral translation as well as yaw rotations of the bees as they approached the gap.

The difference in optic flow across the edge of the wall was normalized with respect to the optic flow 12 mm outside the gap edge, on the wall containing the gap, to reveal the mean motion contrast for the different conditions, see Fig. 5a. For all flights only the flight trajectories of the bees when they were <75mm to the gap were considered for the contrast estimation. A high motion contrast was present only when the distance between the gap and rear wall was large and it monotonically decreased with decreasing distance. Concomitant with the decreasing motion contrast an opposite trend was noted with the time spent by the bees in the near vicinity of the gap (<75mm) before passage, Fig. 5b. Bees spent a longer time <75mm to the gap as the distance between the between the gap and rear wall decreased from 550 to 0 mm, Fig. 5b. For the extreme condition when gap passage was impossible (d = 0) the time spent was not calculated since the bees continued to traverse laterally and no attempts to pass were made. We can conclude that the bees spend more time exploring the situation close to the gap when the optic flow contrast across the gap and, thus, the distance between the gap and the rear wall gets smaller. At the same time, they increase the lateral velocity as a means to increase velocity contrast.

**Figure 5.**
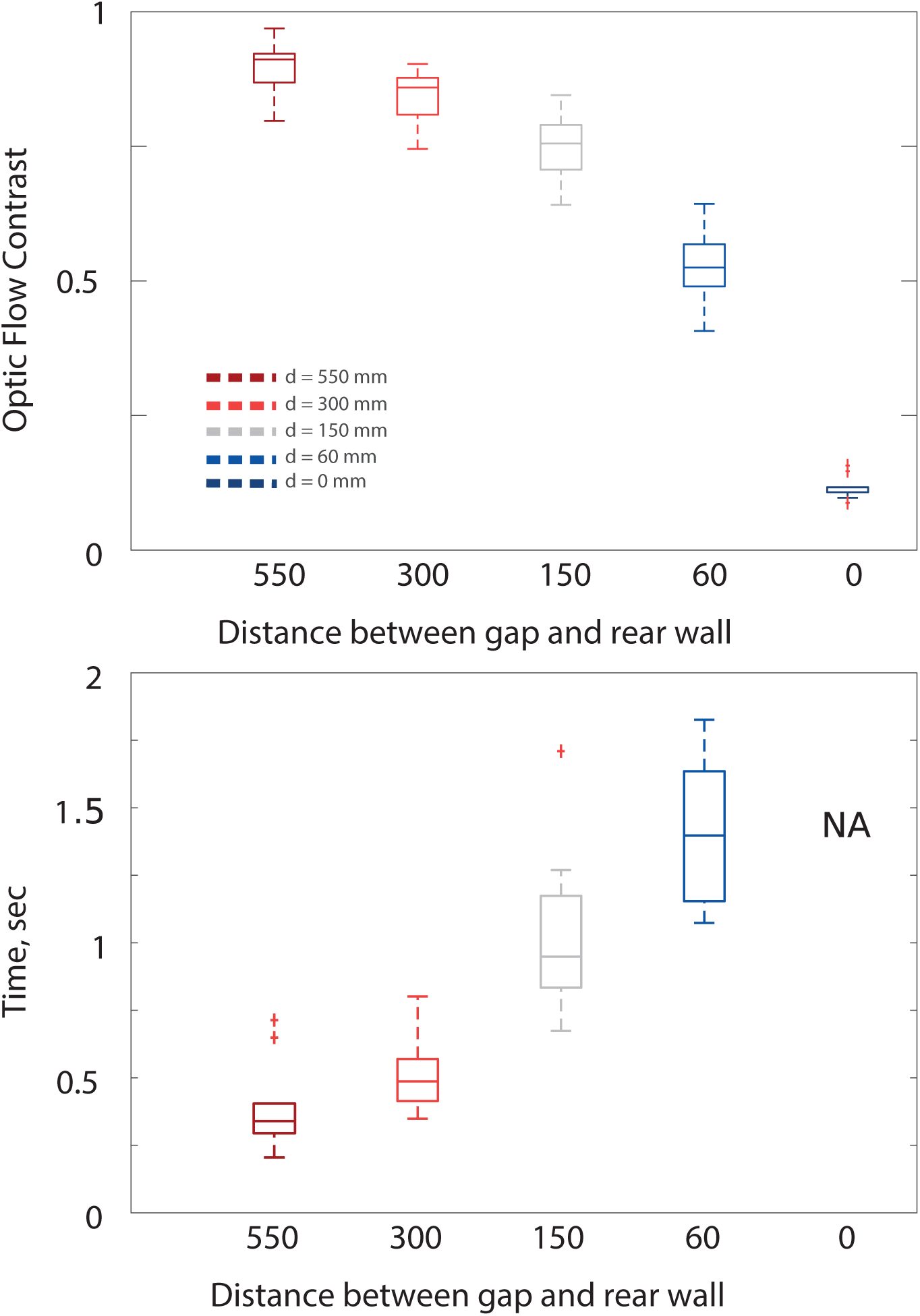
Optic flow contrast measured as the ratio between mean difference in optic flow across the edge of the gap and the optic flow along the outer edge of the gap. (b) Time spent by the bees in the vicinity of the gap (< 100mm) for the different experiment conditions.

## Discussion

Despite their tiny brains, bumblebees and other eusocial insects display a remarkable capacity for navigation through inherently complex environments. The most elemental aspect for locomotion through a cluttered terrain is the identification of a gap between obstacles and subsequently assessing passability. In our experiments we sought insights into the salient mechanisms utilized by bumblebees to identify a gap when presented with an unexpectedly altered environmental situation relative to the conditions of an unobstructed tunnel the bees were familiar with. The bumblebees could thus not learn the gap properties. By considering only bees that were used to returning to the hive through a familiarized unobstructed flight tunnel we exploited the high motivational state of the organism in identifying a route through the altered environment that required passage through the gap. In an alternative setup when the gap was presented to bees that were en route from the hive to the foraging chamber, the bees were much less amenable to the experimental paradigm and chose to return to the safety of the hive.

Bees and other insects might also rely on other cues in facilitating gap detection and passage decision making when flying through cluttered environments such as brightness as shown by (Baird and Dacke, 2016). The few conditions in our experimental analysis where a white background was placed immediately rear of the gap created a scenario where high optic flow difference across the gap was present but nearly zero optic flow within the gap. In this condition none of the bees attempted to pass suggesting that apart from the difference in optic flow across the edges of the gap, a non-zero optic flow within the gap maybe one of the conditions necessary for passage, because only with optic flow inside the gap information about the spatial situation behind the gap may be available. Additionally, the homogenous lighting used in our setup might have not created the necessary brightness difference across the edges of the gap to elicit a brightness-based response. Further investigations are necessary to identify the presence of such virtuosic strategies.

## Approach

When the bees were far from the gap their flight trajectory seemed to be driven by the well-established mechanism of equalizing bilateral optic flow. Since the spatial information on the side walls of the tunnel were similar, nominally consisting of amplitudes varying by 1/frequency (Monteagudo et al., 2017) and a nominally homogeneous illumination, the bees flew close to the centerline of the flight tunnel. This is a familiar feature observed in a number of previous studies that have utilized flight tunnels to study insect and bird flight (Bhagavatula et al., 2011; Schiffner et al., 2014; Srinivasan, 2010). Apart from balancing bilateral optic flow, the flight speed has also been shown to be dependent on the overall width of the flight tunnel as well as its texture. Under the conditions of our experiments, the bees flew at around 0.7m/s in the far field of the gap which was similar to those measured by (Baird et al., 2010) where a similar experiment paradigm was used. Smooth sideward motion interlaced the longitudinal velocity in the bees Fig. 2b-f, this lateral “casting” motion is also a common feature noted in previous experiments on bumblebee flight (Chang et al., 2016; Dyhr and Higgins, 2010; Linander et al., 2015; Ravi et al., 2013).

At around 375mm from the gap, evidence of changes in behavior is first noted as a reduction in flight speed, Fig. 3a. Insects and birds have been shown to respond to unpredictable or unfamiliar situations with reducing speed (Williams and Biewener, 2015). In this region of the flight tunnel the angle subtended by the obstructing wall containing the gap and the gap itself is 36° - 42° and 9° - 14° respectively. (Baird et al., 2010) reported that bumblebees modulate their flight speed using the frontal optic flow and showed that bees responded to abrupt changes in flight tunnel width when it subtended between 23° - 30° on the retina, which is consistent with our data. The bees could either be responding to the obstructing wall, the properties of the gap or the combination of the two. In eliciting a change in flight speed of the bees there appears to be a combined influence of the obstructing wall and the distance between the gap and rear wall when the bees were between 375 – 225mm from the gap. In this region the first consistent reduction in flight speed of the bees across all conditions compared to when they were > 375mm was noted however it was statistically significant only when d< 60mm, see Fig. 2a. Within this region, the mean optic flow difference across the edge of the gap was 60 – 120°/sec when d = 550mm, while the optic flow on the wall along the gap edge was only 3 – 12 °/sec for the extreme non-passable condition (d = 0). The deceleration of the bees for the non-passable condition may be considered as that elicited purely by the obstructing wall. This suggests that in our experimental paradigm the prominence of the gap modulated the approach flight speed of the bees. Comparatively, when they were <225mm from the gap further reduction in flight speed appeared to be mainly influenced by the distance between the gap and rear wall (see, Fig. 3a).

## Gap Perception

For all conditions concomitant with decelerating longitudinal flight speed the bees increased their lateral speed as they neared the gap (Fig. 2). The bees also increased both lateral displacement and speed significantly as the distance between the gap and rear wall was decreased (Figs. 2 & 3). The total accelerations during these lateral maneuvers are higher closer to the gap and mainly oriented normal to the body long axis (Fig. 3). In such cases, the bees are “side slipping” performed by rolling their body to redirect their aerodynamic force vector in the direction of movement, similar to a helicopter (Ravi et al., 2016; Taylor, 2001). Such body roll mediated lateral maneuvers are usually coupled with synchronous counter-rotations of the head to maintain a stable visual field (Boeddeker and Hemmi, 2010; Doussot et al. in prep.). Performing lateral maneuvers where body yaw is limited significantly increases the lateral translational optic flow.

Flies, wasps and a number of other volant insects actively shape the optic flow on the retina by modulating their head and body trajectory to increase the translatory component while minimizing rotations (Egelhaaf et al., 2014). Optic flow derived from translation contains information on the relative distance between environmental features such as obstacles while optic flow from rotations lacks this vital information wherefore volant insects tend to restrict rotations to rapid saccades (Egelhaaf et al., 2012a). Bumblebees flew in our tunnel with minimal yaw rotations (Fig. 1b-f) thus increasing translational optic flow. Optic flow derived through pure longitudinal motion is not much sensitive to distance differences in the frontal visual field, since the flow vectors are small close to focus of expansion. Increased optic flow sensitivity to distance differences between environmental features in the frontal visual field can be achieved, however, through lateral translation (motion parallax) (Collett, 2002). Even when the bees were seemingly uninfluenced by the obstacles (>375mm, Fig. 2), their flight path consisted of smooth lateral movements – casting. The significance of these voluntary lateral movements performed by the bees (Ravi et al., 2016) are unclear. However, they are likely to be used to increase lateral translational optic flow and, thus, to aid depth perception.

A consequence of the increased lateral translations performed by the bees in the proximity of the gap is the large difference in optic flow across the edges of the gap, thereby increasing their salience (Fig. 4b-f). An example of active shaping of optic flow through flight maneuverers can be seen by comparing the generated optic flow when the rear wall was 150mm and 60mm, respectively, behind the gap (Fig. 4a). For these two conditions, when the bees were far (>225mm) the difference in optic flow across the gap edge was significantly higher than when the d > 150mm. However, in near field of the gap (<75mm), due to the increased lateral maneuvers of the bees, the optic flow difference across the gap edge was with 200 – 400 °/sec similar to the d = 550mm condition (Fig. 4a). Active maneuvering in order to discern depth and increase salience of the edges appears to be a compensatory strategy of the bees to the changing distance of the rear wall. Wasps, honeybees and other insects have been observed to perform nominally similar flight maneuvers, which consist of large lateral components, for instance, during their learning flights after leaving an attractive goal location, such as a food source or a nest hole (Zeil, 1996; Dittmar et al. 2010; Lobecke et al. 2018). Here we show that bees actively modulate such behavior in a gap perception context and it appears to depend on the salience of the gap.

In this case bees are likely utilizing a combination of information about the velocity of self-motion, which is related to the input motor signals, and the relative optic flow in discerning the gap salience. The monotonic increase in lateral velocity with decreasing distance between the gap and rear wall does not increase at the same rate until the extreme condition when the gap and rear wall are adjacent (Fig. 2c). Our results suggest that it is likely that there exists a threshold dependent on salience of the gap. We believe this threshold might reflect “passability” based upon identifying gap properties including depth through lateral maneuvering. If the bees cannot assess safe passage, they might resort to searching for alternative gaps in the environment or may fly back. The nominally crescent-shaped flight pattern of the bees close to the gap for the cases where the rear wall was <60mm bears similarity to searching flights and orientation or learning flights performed by bumblebees upon their first departure from their nest hole where they are assumed to probe the layout of the behaviorally relevant nest hole environment (Lobecke et al., 2018; Philippides et al., 2013; Riabinina et al., 2014).

When confronted with a wall unexpectedly blocking their flight path, the bees search for a gap and, therefore, need to probe the spatial layout of the environment. In such conditions bees might utilize a number of cues, including brightness differences, to ascertain the presence of gaps (Baird and Dacke, 2016), but, in particular as shown in the present study, optic flow information. A number of factors are also likely to influence the flight pattern of the bees in this condition including the geometry of the flight tunnel and the obstacles. However, the similarity between the flights when negotiating the gap in our flight tunnel to learning and searching flights of bees observed in the context of local homing behavior merits further investigation. We suggest that in both situation these characteristic meandering lateral flight manoeuvres serve the same basic purpose, i.e. probing the spatial layout of the environment.

## Time to Decision

When flying within a complex cluttered environment an animal constantly needs to evaluate the environmental features confronting it and to make decisions that influence the flight course. Bees spend significant time in the near vicinity of the gap while performing the rapid lateral maneuvers (Fig. 5b). The consistent repeated flights of the bees, especially when the rear wall was <60mm to the gap, suggests that through these flights the bees not only discern the gap geometry but also evaluate passability. Once the potential for safe passage is established traversal through the gap occurs. Measuring the time spent by the bees within the region where most of the lateral maneuvers occur might provide an indication of the time taken by the bees in arriving at a decision. Among all flights recorded none of the bees performed abrupt corrective maneuvers once gap traversal had commenced indicating that decision-making occurs ahead of the gap.

The optic flow contrast appears to be a critical parameter due to a strong and direct inverse relationship with respect to the time spent by the bees evaluating the gap (Fig. 5). Through the repeated lateral movements, the bees appear to establish saliency of and confidence about the geometry by actively generating visual information about passability. The smaller the salience of the gap, the larger are the sideways velocities in order to increase optic flow contrast and the longer do the bees probe the environment to increase their confidence about the situation. As a consequence, decision-making is delayed. Other factors such as familiarity and experience, though unlikely to play a significant role in these experiments, are also likely to influence the bees’ decision time in assessing gap properties and passability in their natural environment, and further experiments are necessary to quantify the influence of these factors on the neural and biophysical mechanics of locomotion through spatially complex environments.

## Acknowledgements

The project is funded by the Alexander von Humboldt Foundation through a Research Fellowship awarded to Sridhar Ravi as well as supported by the Deutsche Forschungsgemeinschaft (DFG grant number OJNB: EG82/19-1) and by the Cluster of Excellence EX 277 Cognitive Interaction Technology (CITEC), funded by DFG.

